# Accessibility of covariance information creates vulnerability in Federated Learning frameworks

**DOI:** 10.1101/2022.10.09.511497

**Authors:** Manuel Huth, Jonas Arruda, Roy Gusinow, Lorenzo Contento, Evelina Tacconelli, Jan Hasenauer

## Abstract

Federated Learning (FL) is gaining traction in various fields as it enables integrative data analysis without sharing sensitive data, such as in healthcare. However, the risk of data leakage caused by malicious attacks must be considered. In this study, we introduce a novel attack algorithm that relies on being able to compute sample means, sample covariances, and construct known linearly independent vectors on the data owner side. We show that these basic functionalities, which are available in several established FL frameworks, are sufficient to reconstruct privacy-protected data. Additionally, the attack algorithm is robust to defense strategies that involve adding random noise. We demonstrate the limitations of existing frameworks and propose potential defense strategies analyzing the implications of using differential privacy. The novel insights presented in this study will aid in the improvement of FL frameworks.

## Main Text

Large-scale data sets have been shown to be highly valuable for data-driven discovery in various fields such as clinical research [1–4], self-driving cars [5, 6], and smartphone keyboard word predictions [7, 8]. The COVID-19 pandemic has highlighted the importance of the rapid acquisition of new evidence for interventions in public health. Yet, data are often collected by different sides, e.g. hospitals, and established legal frameworks limit direct sharing [9], reducing the speed and statistical power of the analyses with possibly harmful consequences for patients [10]. To facilitate the integrative analysis of distributed data sets, federated learning has been introduced by Google Researchers in 2016 [11]. This supposedly allows for privacy-preserving estimation of statistical models from distributed data, making it an essential tool for the rapid assessment of new treatments to improve the fast acquisition of evidence-based interventions in public health. Security is a key topic in the field, as data leakage can result in deontological and consequentialist privacy harms [12].

Federated learning is based on sharing informative summary statistics by individual data owners (each running a data server) with a central hub (Figure 1D). This central hub is responsible for model building. Servers do not share individuals’ data but only non-disclosive summary statistics. This approach is considered privacy-preserving. The focus on privacy in these areas has naturally precipitated extensive research on potential attack vectors. In particular, that sharing parameter gradients – a particular type of summary statistic – in deep neural structures can reveal the training data [13–17]. Algorithms were able to recreate images and texts [16, 17]. Further, data leakage threats have been summarized [18].

**Figure 1:**
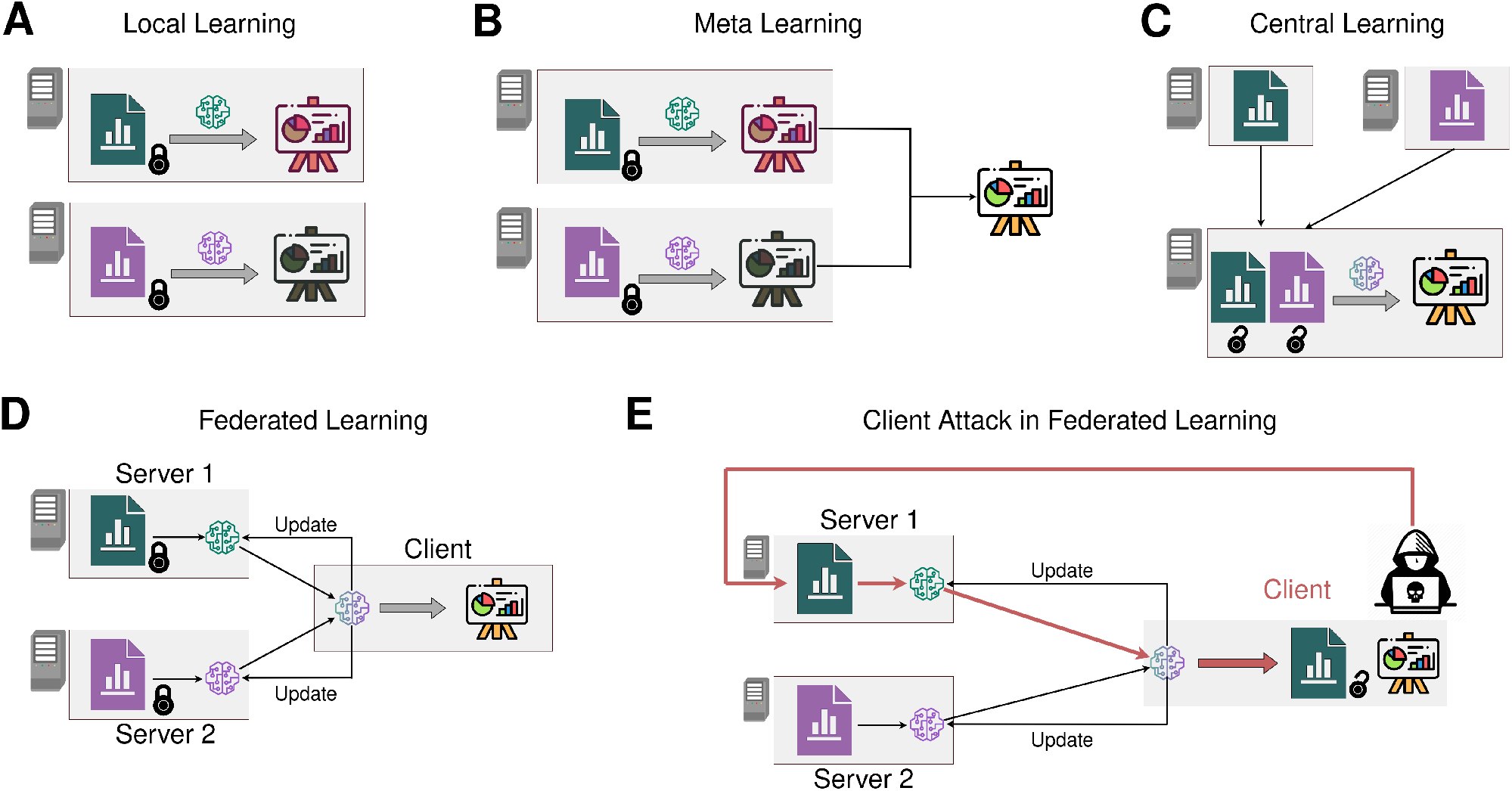
Concept of (attacks in) Federated Learning. **A** In Local Learning, all models are trained separately on different servers. **B** In Meta Learning, all models are trained separately on different servers but individual results are subsequently averaged to obtain meta results. **C** In Central Learning, the data is pooled and one model is trained. Hence the data must be shared. **D** In Federated Learning, the data is kept private on the servers. One model is trained with continuous updates between the client and the servers. **E** Illustration of a client side attack in Federated Learning. A malicious client uses the information received from the server to retrieve private data. This figure has been designed using resources from Flaticon.com.

In this study, we complement the previous work by focusing on attacks based on basic functionalities that are available in established Federated Learning frameworks. We consider the possibilities of a malicious client who tries to obtain the data stored across different data owners and introduce a new attack concept. To perform the attack, we generate known linearly independent vectors on each server. After concatenating them on the client side, we use sample means and sample covariances to reconstruct the server side data nearly up to numerical precision (Fig 2). In contrast to the well-studied gradient approach, the presented method requires comparatively little time, and no model knowledge. Moreover, our algorithm carries desirable statistical properties: Repeated execution allows for exact data reconstruction even if random additive noise is applied to the covariance and means. In our opinion, this combination of features makes this attack strategy more problematic than any approach outlined previously. We discuss our algorithm theoretically and demonstrate its use in the open-source frameworks R DataSHIELD (version 6.2.0) [19], TensorFlow Federated (version 0.36.0) [20] and implemented a proof-of-concept for further platforms using PyTorch [21].

**Figure 2:**
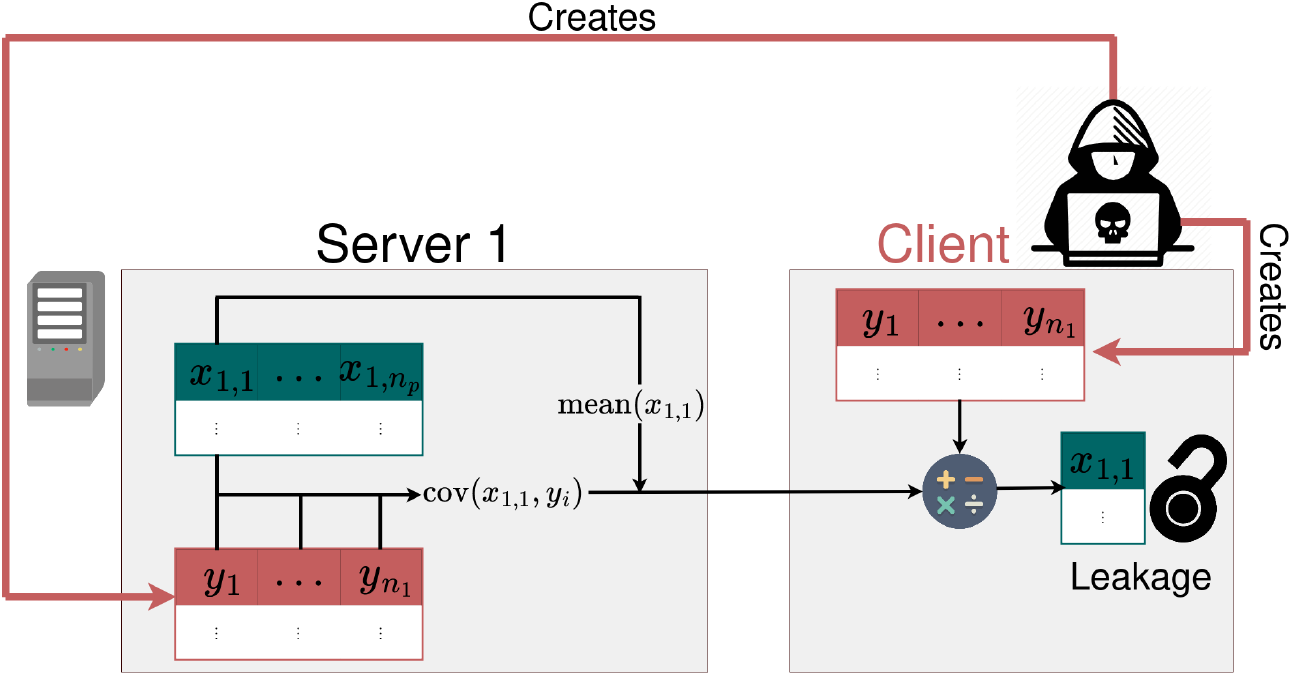
Covariance-Based Attack Algorithm setup for reconstructing data *x*_1,1_ on the first server. The malicious client generates linear independent vectors 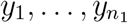 on the server and client side, computes the covariances of them together with the attacked vector *x*_1,1_, and returns them with the mean of *x*_1,1_ to the client side. Subsequently, the returned information is used with the means of 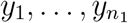 to compute *x*_1,1_ on the client side. The algorithm can be repeated for all *x*_*j,k*_ to obtain the full data set. This figure has been designed using resources from Flaticon.com.

### Covariance-Based Attack Algorithm

The distributed infrastructure consists of *n*_*h*_ servers. The *j*-th server hosts observations *s* = 1, …, *n*_*j*_ (e.g. patient data sets), each with information for variables *k* = 1, …, *n*_*p*_. Accordingly, each server stores a data matrix 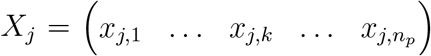 with 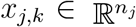, where each vector *x*_*j,k*_ contains information about the variable *k* for all *n*_*j*_ samples on the server *j*. Without loss of generality, the malicious client focuses on a specific variable *k* on a specific server *j*, denoted by *x*_*j,k*_. Retrieving the remaining variables and servers can be subsequently obtained analogously. We assume that the attacker has at least three basic tools: (T1) a sample mean function Mean(*x*), (T2) a sample covariance function Cov(*x, y*), and (T3) an algorithm 𝒜 generating *n*_*j*_ linearly independent vectors 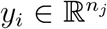 on the server side and their column-wise collection as a matrix 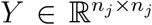 on the client side, 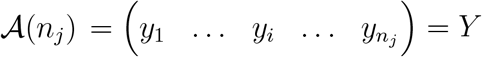 with 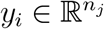 (Figure 2).

These requirements are met by many distributed analysis frameworks, virtually all of which include functions for computing sample means (T1) and covariances (T2). The input of the covariance function is usually not restricted to subsets of the data matrix X but allows for other inputs *y* (T2). The availability of a function for the construction and sharing of linearly independent vectors (T3) might seem the least obvious, but it is available in most tools. For instance, it is necessary in the context of optimisation via federated averaging: The client receives the server side gradients, updates the parameters, and sends them back to the servers. Therefore, for any system where this operation is possible, assumption (T3) must be satisfied, since we need to be able to sent vectors from the client side to the server side. Well-known and widely used distributed analysis frameworks for which assumptions (T1)—(T3) are fulfilled are TensorFlow Federated [20] and DataSHIELD [19].

Assuming that (T1)—(T3) are met, the centerpiece of our algorithm is the fact that evaluating the sample covariance makes it possible to reconstruct the inner vector products between the attacked vector *x*_*j,k*_ and the linearly independent vectors 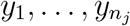. This yields a linear system of *n*_*j*_ equations which can be solved for *x*_*j,k*_ and written in matrix form as

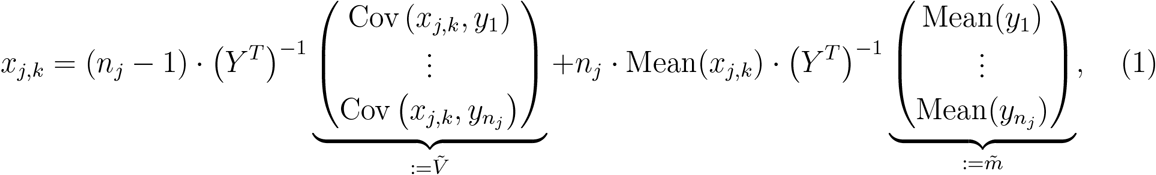

where the right-hand side of (1) is known by the malicious client. Derivations of the computations are reported in the section Mathematical Computations of the Methods. The presented procedure can be repeated for each variable *k* = 1, 2, …, *n*_*p*_ and server *j* = 1, 2, …, *n*_*S*_, using the same or newly generated linearly independent vectors *y*_*i*_, until all data 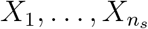 is obtained. As the covariance calculation is essential, we refer to the strategy as *Covariance-Based Attack Algorithm*.

#### Algorithm 1 Covariance-Based Attack Algorithm

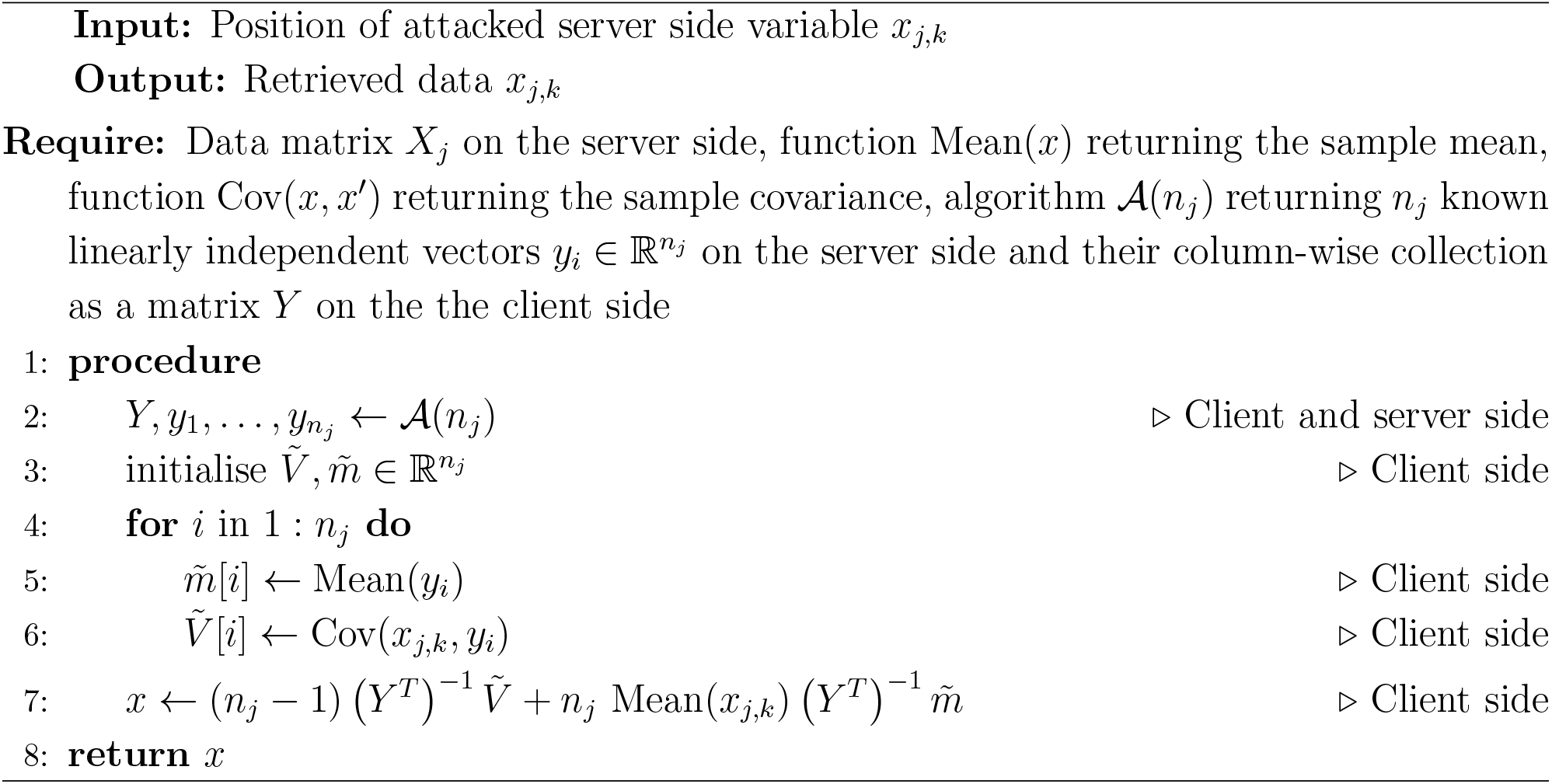

### DataSHIELD is vulnerable to the algorithm

To demonstrate the Covariance-Based Attack Algorithm and the vulnerability of existing distributed analysis frameworks, we considered different software packages. First, we provide an example implementation in DataSHIELD (version 6.2.0) and its base package dsBaseClient [19]. This tool is well established and used in various biomedical applications [3, 22–25] in which data sharing is limited, e.g. to ensure compliance with privacy regulations, such as the General Data Protection Regulation (GDPR). The data set utilized is the CNSIM data set from the DataSHIELD tutorial [26] to ensure an easy-to-reproduce test case. This data set consists of 3 servers with a total of 9,379 synthetic observations of 11 personalised obesity-related variables. We have reconstructed information on individual body mass index (BMI) measurements from the first server using a complete case analysis with a sample size of *n*_*j*_ = 250.

DataSHIELD meets the requirements (T1)—(T3) and is therefore vulnerable to the Covariance-Based Attack Algorithm. The functions to compute sample means (T1) and sample covariances (T2) are `ds.mean` and `ds.cov`, respectively. These functions return the means and covariances directly, but require mild conditions on the attacked data *x*_*j,k*_: (C1) the sample sizes *n*_*j*_ must exceed the thresholds *n*_*j*_ > 3 (`ds.mean`) and *n*_*j*_ > 6 (`ds.cov`); and (C2) both levels of a dichotomous variable must occur at least 3 times in the given data vectors. The conditions (C1) and (C2) ensure a privacy-preserving analysis if the functions are applied once. If any of the assumptions (C1) and (C2) were violated, descriptive statistics or further analysis with *x*_*j,k*_ would be impossible. Therefore, it is reasonable to assume their validity. In our example, the data have *n*_*j*_ = 250 observations of a continuous variable so that requirements (C1) and (C2) are clearly satisfied. The construction of *n*_*j*_ linearly independent vectors (T3) can be implemented in several ways. We used the function `ds.dmtC2S` to send client side matrices to the server side. Hence, it is possible for the client to create suitable linearly independent vectors *y*_*i*_ on the client side and to send them to the server side. Note that since the covariance operation is performed on *x*_*j,k*_ and all *y*_*i*_, (C1) and (C2) must hold for all *y*_*i*_ as well. Since *x*_*j,k*_ and *y*_*i*_ have the same length *n*_*j*_, (C1) holds. To meet (C2) and the linear independence condition, we draw each element *y*_*i*_ from a standard normal distribution, so that *y*_*i*_ almost surely consists of *n*_*j*_ distinct entries and that 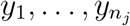 care almost surely linearly independent. In principle, however, the malicious client can use any linearly independent vectors 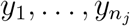 that meet the requirements (C1) and (C2).

After creating all linearly independent *y*_*i*_ on the server and client side and computing the relevant means and covariances, the data can be obtained as described in (1). Our evaluation of the above example shows that the true data can be reconstructed almost perfectly (see Figure 3A). The Pearson correlation coefficient between true and retrieved BMI values is 1.0. The highest absolute error observed is 2 · 10^−12^, which is close to numerical accuracy. This demonstrates that the Covariance-Based Attack Algorithm is not limited to theoretical settings. Instead, data leakage can also be achieved in real-life setups.

**Figure 3:**
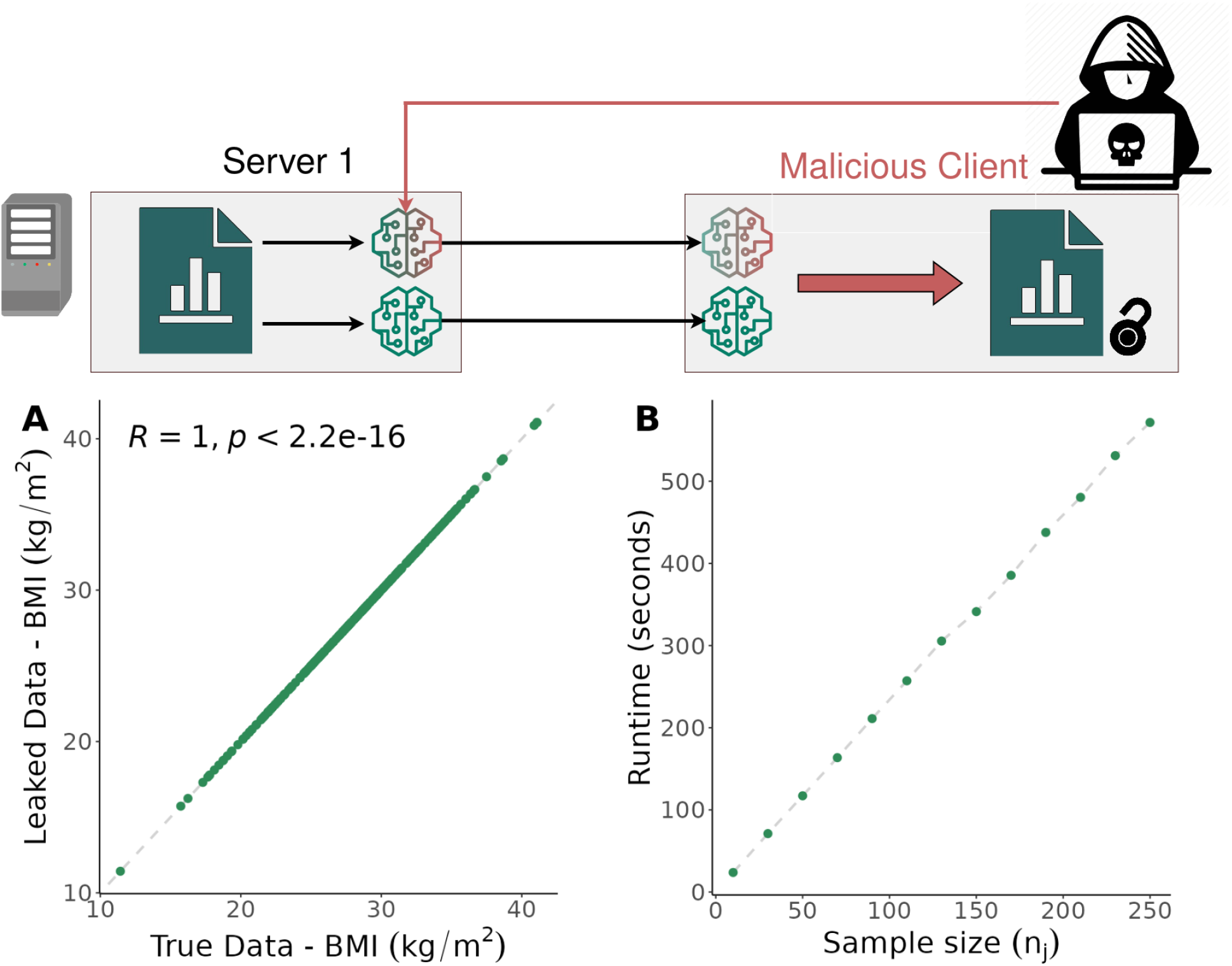
Leakage results and computation times for DataSHIELD. **A** True data values from the first server of the CNSIM data set vs. the corresponding retrieved data provided by the Covariance-Based Attack Algorithm. **B** Computation time of the algorithm for different sample size. This figure has been designed using resources from Flaticon.com.

### Computation complexity of data reconstruction grows linearly with sample size

To study the applicability of the Covariance-Based Attack Algorithm, we considered the scaling of the computation time with growing sample size *n*_*j*_. As computation time, we consider the wall time required to obtain the result.

In theory, the sample size determines the time requirements in different ways. First, it determines the size of the system of equations (1). This size is identical to *n*_*j*_, meaning that *n*_*j*_ requests must be sent to the *j*-th server. The communication overhead for a request is constant, but the computation time will generally grow linearly with *n*_*j*_ [𝒪(*n*_*j*_)], as the dimensionality of the scalar product increases. Secondly, the computation time for solving the linear system from (1) grows cubically 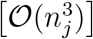 using the `solve` command in R and the `linalg.inv` command in Python available through the NumPy library. Hence, there are linear, and cubic contributions, with different pre-factors, to the computation time.

To evaluate the scaling behavior in practice, we considered subsets of the CNSIM data set of different sizes and determined the wall time required to complete the attack (Figure 3B). We observed linear scaling (Figure 3B), meaning that the communication overhead determines overall wall time. Indeed, even for the largest data set considered, matrix inversion required only 0.004 seconds, meaning that it contributed only 7 · 10^−6^ percent to the overall time.

The essentially linear scaling behavior in the relevant regime, compared to the theoretically cubic scaling behavior, leads to this attack being feasible in many real-world scenarios.

### The Covariance-Based Attack Algorithm is robust against noise perturbations in DataSHIELD

The Covariance-Based Attack Algorithm allows for the reconstruction of the data on the servers. We further investigated whether our approach is robust to adding zero-mean noise to the means and covariances before returning them to the client. In this case, the client observes noise-corrupted data estimates

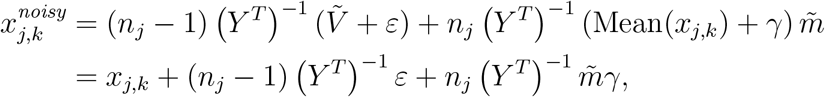

with zero-mean and finite-variance noise terms *ε* and *γ*.

The noise-corrupted data estimate 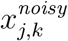 can be decomposed into the true data *x*_*j,k*_ and a noise component so that the malicious client cannot retrieve the original data (Figure 4A). However, the malicious client is, given suitable communication and computational budgets, which is given for example in DataSHIELD, able to run the algorithm *R* times. If *R* is sufficiently large, the zero-mean noise components average out such that the mean 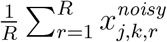 converges in probability to the data *x*_*j,k*_ (Figure 4B). We provide a proof in the Method section. Hence, even if noise is added to the means and covariances, a malicious client can retrieve the data if no further restrictions on the analyst are imposed.

**Figure 4:**
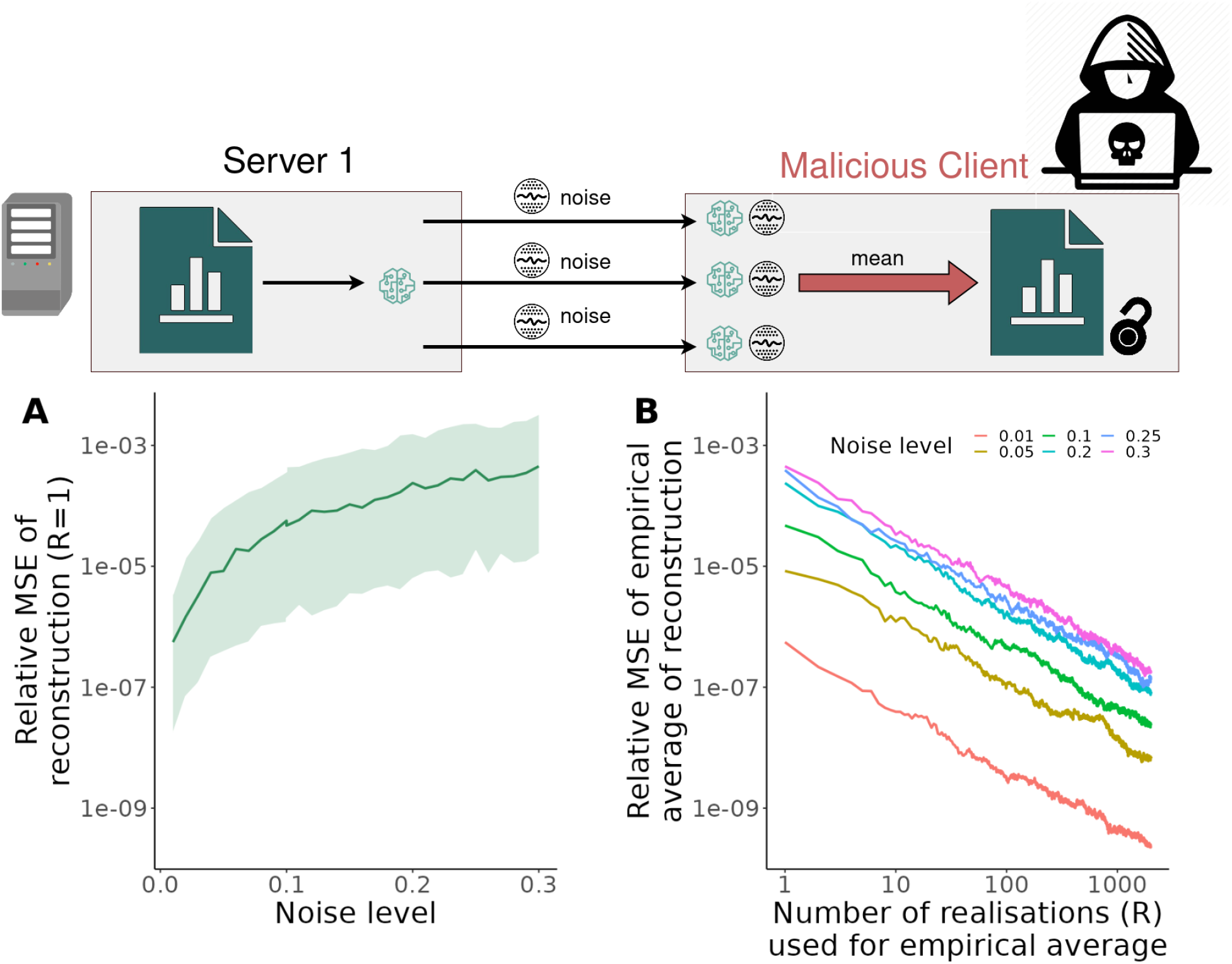
Robustness of Covariance-Based Attack Algorithm to normally distributed noise on means and covariances. **A** Relative mean squared error (MSE) of the reconstructed data values for different noise level if only a single realization is available (*R* = 1). The median (line) and the 5th to 95th-percentile (area) of 200 replicates are depicted. **B** Relative mean squared error (MSE) of the empirical mean of reconstructed data values obtained from different numbers of realization (*R* = 1, …, 1000) and four different noise levels. The median (line) of 200 replicates is depicted. This figure has been designed using resources from Flaticon.com.

### TensorFlow Federated is vulnerable to the algorithm

To assess whether other tools allow for the implementation of similar attack strategies, we considered TensorFlow Federated (version 0.36.0)[20]. In contrast to DataSHIELD, this open-source framework for computations on decentralized data is meant for experimentation with Federated Learning. Yet, if experimentation environments allow for (non-trivial) disclosive computations, these are likely to find their way into application. Accordingly, we evaluated the possibility of implementing the Covariance-Based Attack Algorithm using a set of basic functions.^1^

Our assessment revealed that TensorFlow Federated meets the tool requirements (T1)—(T3) and allows for the implementation of the Covariance-Based Attack Algorithm. Functions can be constructed in TensorFlow Federated by wrapping functionalities from Python packages, e.g. TensorFlow or numpy, in a function and labelling it with `tf computation`. To compute sample means (T1), a function that computes the average of *x*_*j,k*_, e.g. using `numpy.mean`, can be implemented. For (T2), one can for instance wrap the function `stats.covariance` from the TensorFlow probability package. Both functions need to be applied with the functionality of `federated map` to return values from the server side. Since TensorFlow Federated does not enforce further privacy leakage checks, these functions do not have requirements that are equivalent to (C1) and (C2) for DataSHIELD. However, we expect that if TensorFlow Federated is used in real-world applications, further disclosiveness checks, similar to (C1) and (C2), will be implemented. For (T3), TensorFlow Federated offers the `tff.federated broadcast` function which is similar to the function `ds.dmtC2S` as it sends objects from the client to the server side. Due to the current lack of requirements such as (C1) and (C2), the vectors 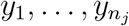 must be linearly independent but no further restrictions have to be imposed.

The implementation of the Covariance-Based Attack Algorithm in TensorFlow Federated was applied to the aforementioned CNSIM data set. We found that this allows for a reconstruction of the data up to numerical accuracy (Supplementary Figure 6). Hence, data leakage is also possible in TensorFlow Federated, using algorithms that appear to be non-disclosive. This raises questions regarding the suitability of the framework for experimentation with Federated Learning.

### PyTorch based platforms used in real-world applications are vulnerable to the algorithm

Further Federated Learning tools, such as Nvidia Flare [27], IBM Federated Learning [28], Intel’s OpenFL [29], and PySyft [30], are widely used in real-world applications that handle sensitive data. For example, Nvidia Flare is used by Microsoft Azure, American College of Radiology, Rhino Health and others as indicated on their website.^2^ These tools advertise their flexibility in using popular Python-based deep learning packages like PyTorch [21]. In light of the potential for malicious implementation of algorithms, we evaluated the feasibility of implementing the Covariance-Based Attack Algorithm on these platforms.

Our analysis demonstrated that the platforms under examination fulfill the necessary tool requirements (T1)–(T3), enabling the implementation of a Covariance-Based Attack algorithm. Users of these platforms have the ability to specify their models in PyTorch, Tensorflow, or Numpy, and the platforms subsequently integrate the specified models into a federated learning workflow so that the final model specification is ultimately determined by the client. To compute sample means (T1) one can apply `torch.mean` on the variables of interest. Regarding tool requirement (T2), one could use the PyTorch function `torch.cov`. However, no direct algorithm for generating linear independent vectors (T3) is available. However, linear regression models `torch.nn.Linear` that incorporate the attacked variable as the dependent variable and the randomly generated vectors (T3) as independent variables, since users have full control over specifying any type of model in PyTorch. Hence, one can easily specify multiple linear models, thus incorporating the generation of (T3) in the specification of the model. The least squares estimate of *β* for the model *x*_*j,k*_ = *y*_*i*_*β* + *ε* can be represented as 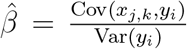. By rearranging this expression, we can express the covariance as 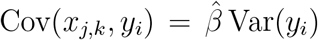, where 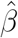 is obtained through linear regression and Var(*y*_*i*_) is calculated easily on the client side.

The examined platforms do not enforce (C1) and (C2) as built-in security measures. However, additional security features are implemented, for example, models must undergo a review process prior to being utilized with real-world data. Our work can improve this process by raising the awareness to potential disclosive models. Moreover, the execution of algorithms might be limited by a privacy budget in accordance with differential privacy principles, as outlined in [31].

### Differential privacy limits the algorithms use but precludes standard analysis

Our proposed algorithm, the Covariance-Based Attack, demonstrates effective data retrieval from platforms that do not implement computational privacy budgets. To assess the impact of such budgets on our method, we evaluated its robustness by testing its performance when platforms enforce differential privacy restrictions on the client’s analysis. Differential privacy improves security by adding appropriate noise to the output of an algorithm, 𝒜, when applied to a data set, 𝒟, in order to ensure that no individual in 𝒟 can be directly identified [31].

The privacy budget, *ϵ*_𝒜_, which controls the level of noise added, is allocated to the analyst and must be used with caution, as it limits the number of algorithm calls that can be made by the analyst.

The privacy budget consumed by the Covariance-Based Attack algorithm, denoted as *ϵ*_att_, can be upper-bounded by the sum of the individual privacy budgets allocated for the computation of covariances and means

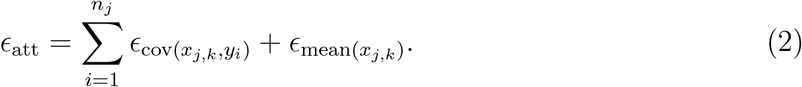

Detailed mathematical descriptions of the privacy budget consumption of the mean 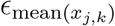 and covariances 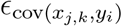 are given in the Methods section.

By specifying a privacy budget, data owners face the trade-off between more function calls or increasing security. More function calls could result in more informative outcomes but increases the risk of data leakage, whereas less function calls increase security but could limit the statistical results. We examined this trade-off using the CNSIM data set. First, we computed the possible number of runs of the Covariance-Based Attack Algorithm depending on the privacy budget, and examined how the relative error of reconstruction changes with the privacy budget (Figure 5A,B). We varied the privacy budget parameter, *ϵ*, between ln(1.01) and ln(3), based on the range defined in [31]. Second, we computed covariances (Figure 5C) between variables of the CNSIM data to see how much privacy budget is consumed by standard descriptive statistics. Third, we analyzed the consumption of the privacy budget for a basic analysis workflow with means, and (Co-)Variances of two variables (Figure 5D) examining if a standard analysis is still feasible under the restrictions of differential privacy.

**Figure 5:**
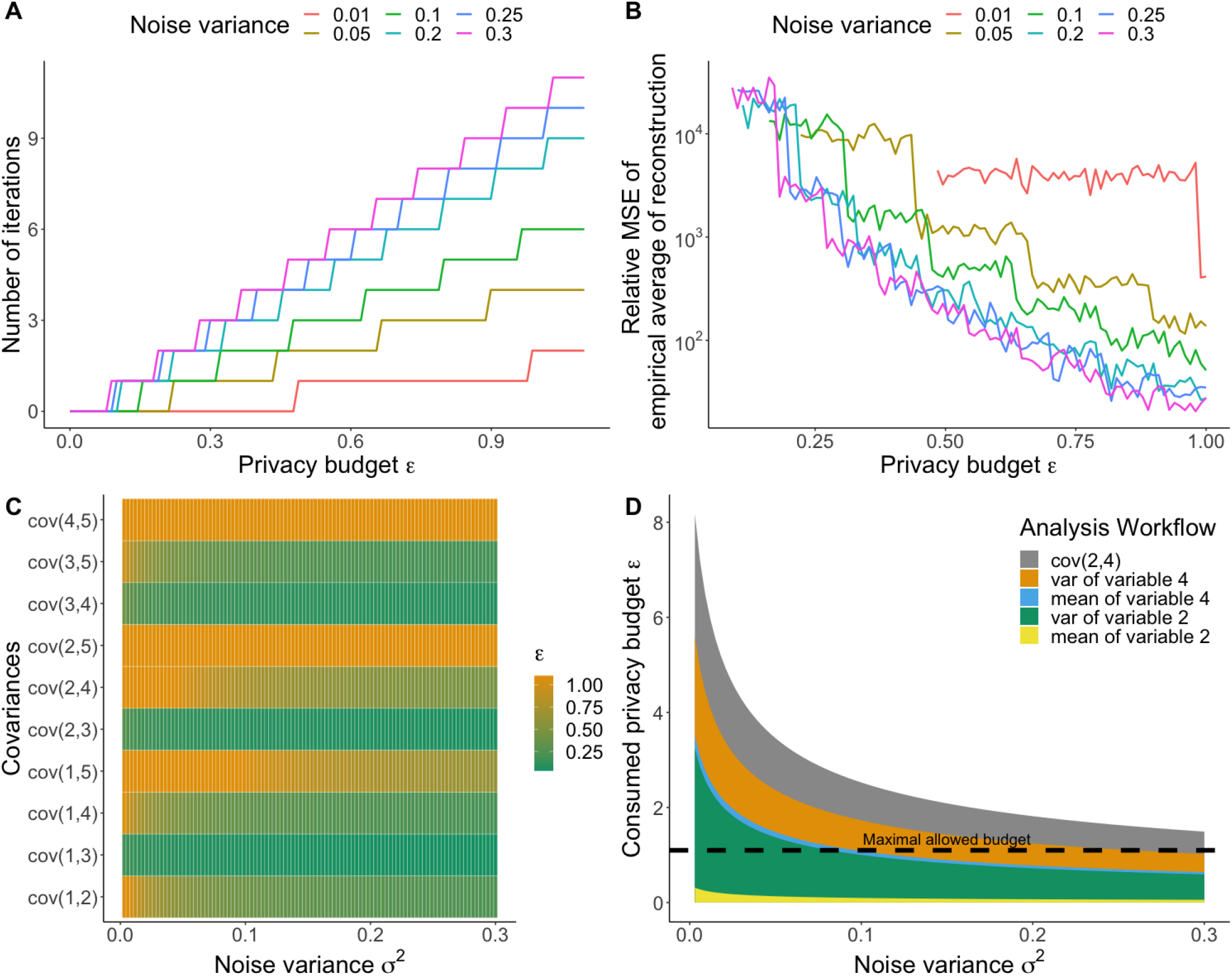
**A** Number of possible iterations of the Covariance-Based Attack Algorithm for different noise levels still satisfying *ϵ*-differential privacy using the CNSIM data set. **B** Error of the Covariance-Based Attack Algorithm after allowed numbers of iteration for different noise levels still satisfying *ϵ*-differential privacy using the CNSIM data set. The median (line) of 200 replicates is depicted. **C** Privacy budget consumed by computing the covariances between metric variables in the CNSIM data set for different noise levels. The consumed budget *ϵ* is truncated at the maximal allowed privacy budget ln(3). **D** The privacy budget consumed by performing a standard analysis on two variables (computing the mean and variance of two metric variables and their covariance).

Increasing the noise variance of the outputs of the covariances and means enables the analyst to perform more iterations of the Covariance-Based Attack algorithm, as shown in Figure 5A. However, even at the highest noise variance, it is not possible to obtain a sufficient number of runs of the algorithm to recover the true data by averaging out the noise, as demonstrated in Figure 5B. Interestingly, more noise leads in general to a higher accuracy of the algorithm for the same privacy budget *ϵ* since more iterations of the algorithm can compensate increasing noise as one can see in Figure 5B.

However, the allocated privacy budgets are not sufficient for a non-malicious analyst to perform covariance computations on the five metric variables of the CNSIM dataset. Even when using high noise variances, the computation of one covariance consumes already the whole privacy budget, as shown in Figure 5C. The standard analysis workflow could not be computed for all considered noise variances due to the consumed accumulated privacy budget, as illustrated in Figure 5D.

## Discussion

Federated Learning has demonstrated its utility and importance in various fields, and its significance has been further highlighted during the SARS-CoV-2 pandemic, with many consortia utilizing it extensively [1, 32, 33]. However, it is crucial to ensure that the data of the participating servers remains protected. To accomplish this, a thorough examination of attack strategies is necessary. In this study, we introduce the novel Covariance-Based Attack Algorithm, which highlights the vulnerability of established Federated Learning systems.

We have demonstrated that a malicious client could leverage the Covariance-Based Attack Algorithm to extract data from a Federated Learning system. Our approach differs from previous studies that focused on information leakage through gradients obtained from deep neural models. It relies on constructing linearly independent vectors on the server side and utilizing sample means and sample covariance functions that are accessible to the client. The attack approach is efficient, scalable, and can be easily implemented, and it is not hindered by noise perturbations if the number of function calls is not restricted. However, implementing differential privacy can prevent the algorithm but would also hinder the application of (linear) analysis tools making differential privacy impractical to use if rather standard tools like linear models should be used. Our algorithm has been successfully implemented on DataSHIELD (version 6.2.0) and TensorFlow Federated (version 0.36.0), where we were able to reconstruct the data. We have informed the respective developers. Furthermore, we have implemented a proof-of-concept using PyTorch (version 1.13.0) showing the usability of the algorithm on other Federated Learning platforms.

Our findings reveal that the existing functionalities of Federated Learning frameworks need to be scrutinized in regards to data leakage threats. To address this vulnerability, we propose to tackle the issue induced by (T3). An effective solution would be to immediately process these vectors within a function instead of creating a vector on the server side avoiding the possibility of computations with client-known objects.

It is important to note that the proposed strategy can only be executed if the attacker can act as the client, which typically requires obtaining log-in credentials. However, even with this access, the security of the data should not solely rely on the trustworthiness of the client, as the Covariance-Based Attack Algorithm demonstrates that data can still be retrieved. The need for Federated Learning would be eliminated if the security of the data is dependent on the trustworthiness of the client alone, as it could be replaced by Centralized Learning. Furthermore, this raises concerns about compliance with the General Data Protection Regulation (GDPR) and about responsibility and liability in the event of unknown attack strategies.

With this work, we contribute to the growing literature on data leakage problems in Federated Learning systems and aim to support research on Federated Algorithms and raise awareness about potential security risks. We did not study other distributed frameworks such as Swarm Learning, but we encourage a thorough review. Although security levels may appear to be higher as the aggregation of information is shared, an attack may still be possible if requirements (T1)—(T3) are met.

We anticipate that our results will play a role in developing design criteria for the architecture of Federated Learning platforms. We have shown that existing systems need to be enhanced to minimize the risk of data leaks.

*Supplementary Information (code) is available for this paper. Correspondence and requests for materials should be addressed to Jan Hasenauer*.

## Methods

### Proof of correctness for the Covariance-Based Attack Algorithm

In this study, we consider an attack by a malicious client and provide an algorithm for data reconstruction based on covariance information. In the following, we provide the mathematical derivation of the algorithm.

For the attack, it is necessary to compute sample means

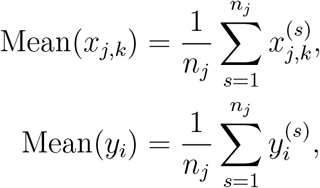

and sample covariances,

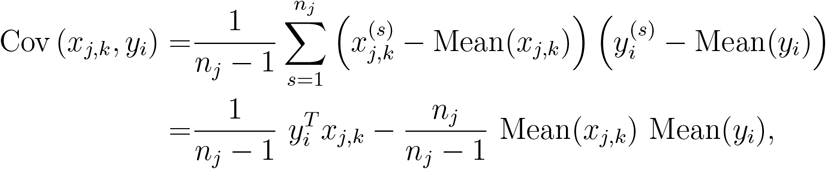

for all *i* = 1, 2, …, *n*_*j*_ on the server side and return them to the client side. The vectors *y*_*i*_ are chosen in a way to ensure their linear independence.

To reconstruct *x*_*j,k*_, we exploit the fact that the sample covariances can be reformulated to determine the inner product 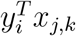,

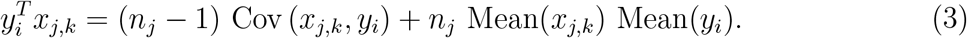

We combine the equations for *i* = 1, …, *n*_*j*_ from (3) to a system of equations in matrix form:

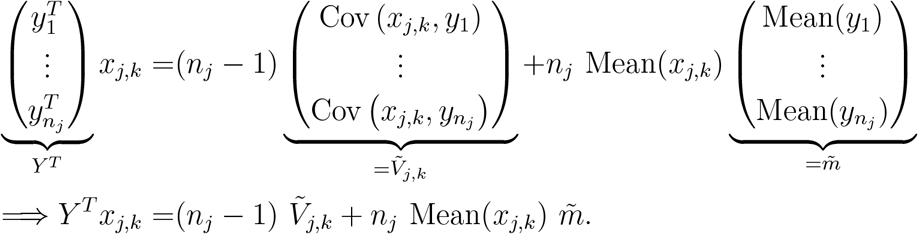

Since the vectors *y*_*i*_ were chosen to be linearly independent, *Y*, and therefore also *Y* ^*T*^, are invertible. Hence, we can multiply both sides of equation (4) by the inverse of *Y* ^*T*^ to obtain

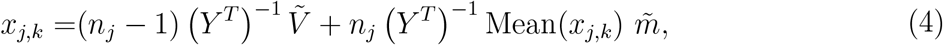

where the right-hand side is known to the client. This provides a constructive proof for the recovery of *x*_*j,k*_ using the proposed approach.

The same procedure can be repeated for all *n*_*j*_ servers and all *n*_*p*_ variables, yielding comprehensive information about potentially sensitive data on the servers.

### Robustness of the Covariance-Based Attack Algorithm to noise perturbations in DataSHIELD

As a defense strategy against malicious clients, we consider the perturbation of mean and covariance with noise but without the additional use of differential privacy. More specifically, we consider the addition of zero-mean noise to means and covariances on the server side before sending them to the client side. Given only access to noisy data, one might assume that the client will not be able to reconstruct *x*_*j,k*_ exactly. However, running the attack algorithm multiple times on the same variable and averaging these results yields a random variable that converges in probability to *x*_*j,k*_ such that the malicious client is, given an appropriate communication and computational budget, able to retrieve all information about *x*_*j,k*_. We prove that the empirical mean of the noisy results of the Covariance-Based Attack Algorithm 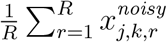, with *R* denoting the number of calls in an attack, converges in probability to *x*_*j,k*_, i.e., formally for any *c* > 0

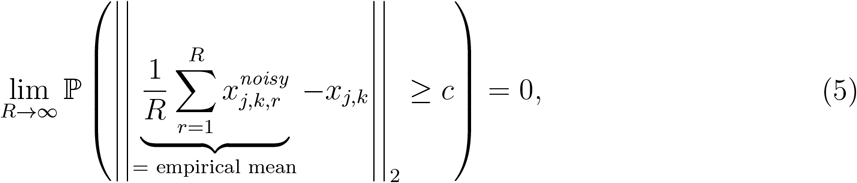

where 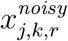 is the result of the *r*-th run of the Covariance-Based Attack Algorithm.

Let *ε*_*r*_ be an *n*_*j*_ dimensional random vector with mean 𝔼(*ε*_*r*_) = 0 and covariance matrix m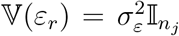 for which 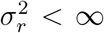. Let *γ*_*r*_ be a random variable with mean 𝔼(*γ*_*r*_) = 0 and variance 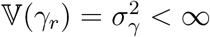. Furthermore, let *γ*_*r*_ and *ε*_*r*_ be uncorrelated so that 𝔼(*γ*_*r*_ · *ε*_*r*_) = 0.

The noisy version of equation (4) is given by

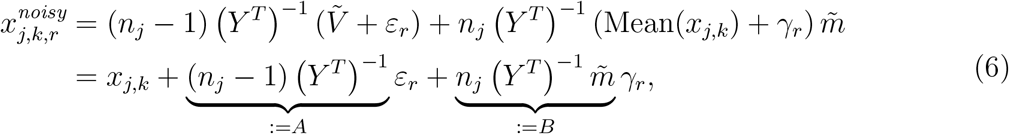

such that 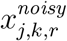 can be decomposed into the true *x*_*j,k*_ and a noise term. Combining (5) and (6) shows that (5) is proven if the mean of the noise term converges in probability to zero, such that it is sufficient to show that

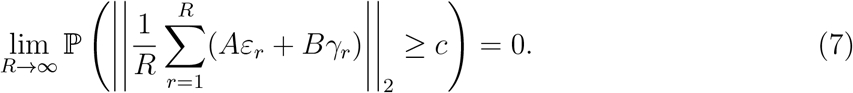

This can be shown by applying Markov’s Inequality

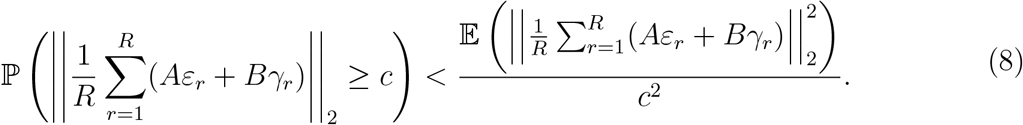

Since (8) holds for all *R*, it is sufficient to show that the numerator of the right-hand side converges to 0 if *R* → ∞ in order to prove (7). To facilitate notation, the entries of *A*^*T*^*A* are denoted by *a*^(*s,s*′)^ and the entries of *ε*_*r*_ by 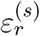. Note that the following holds:

- 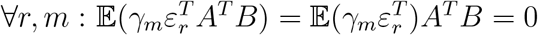,
- ∀*l* ≠ *m* : by independence of *ε*_*r*_ and *ε*_*m*_, 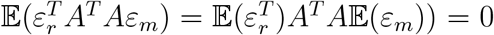 and by independence of *γ*_*r*_ and *γ*_*m*_ that 𝔼(*γ*_*r*_*γ*_*m*_*B*^*T*^ *B*) = 𝔼(*γ*_*r*_)𝔼(*γ*_*m*_)*B*^*T*^ *B* = 0,
- 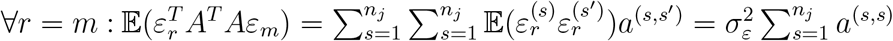 and 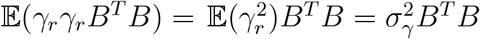.

The numerator of the right-hand side of (8) can therefore be written as

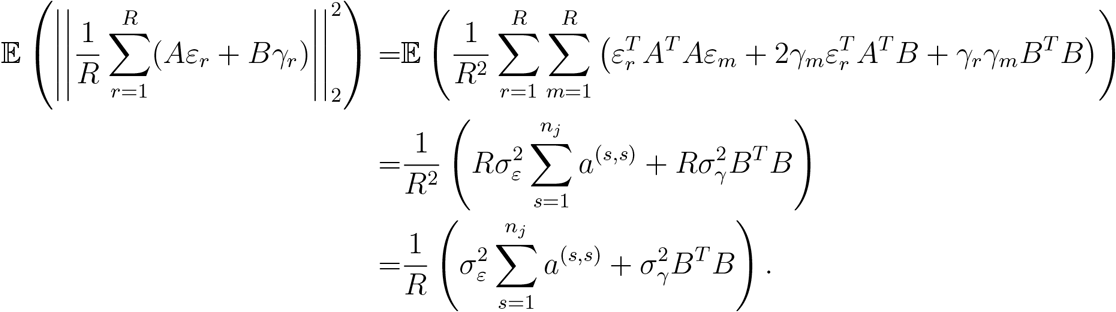

This is a constant multiplied by *R*^−1^. Accordingly, (7) holds and therefore (5) is proven.

In the manuscript, we provide an analysis of the mean squared error for different numbers of calls from an attacker and different levels of noise. The relative mean squared error (RMSE) is here defined as

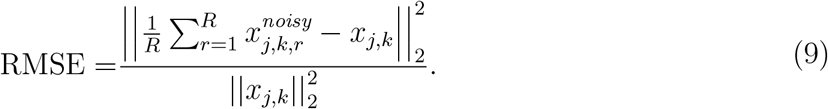

### Derivation of privacy budget consumption of mean and covariances

The privacy budget consumption of the mean (covariance) algorithm is given by

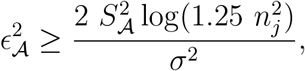

where σ^2^ is the noise level of the normally distributed noise that is added to the output of the mean (covariance). *S*_𝒜_ is the algorithm dependent local sensitivity which is the maximum amount that the output of a function can change given a change of a single input. It therefore determines the amount of noise that needs to be added to the output of a function in order to ensure that it satisfies the privacy guarantees of differential privacy [31]. Mathematically, the local sensitivity *S*_𝒜_ is defined as

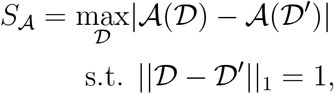

where the L1 norm of a data set indicates how many individuals have been added or removed from a data set. To compute the local sensitivity *S*_mean(z)_ of the mean of a vector *z* = (*z*_1_ … *z*_*n*_)′ ∈ ℝ^*n*^, we denote the maximum absolute deviation of any component of *z* from the mean by *z*\*

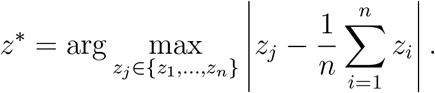

The data set 𝒟 equals the vector *z* and removing one entry from *z* or adding an entry to *z* yields the neighboring data set 𝒟′. The local sensitivity of the mean becomes therefore

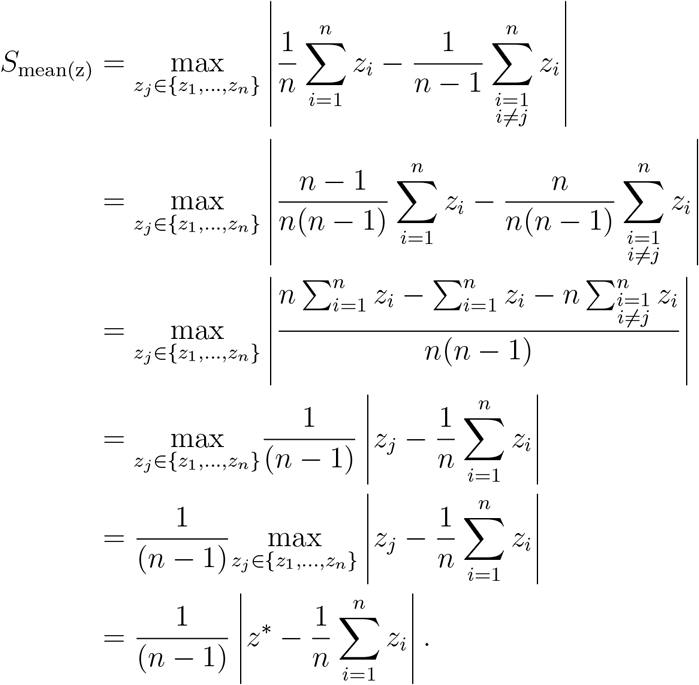

For the local sensitivity *S*_cov(w,z)_ of the covariances for *z* and another vector *w* = (*w*_1_ … *w*_*n*_)′ ∈ ℝ^*n*^, the data 𝒟 equals the matrix (*w z*). The optimization problem becomes therefore

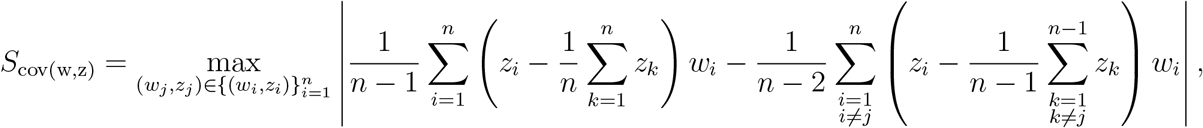

which we solve in our simulation study by evaluating all possible outcomes of the absolute value when removing any (*w*_*j*_, *z*_*j*_).

### Implementation and code availability

A code example of our attack algorithm using the open source frameworks DataSHIELD (version 6.2.0) and TensorFlow Federated (version 0.36.0) together with the proof-of-concept using PyTorch (version 1.13.0) and the differential privacy analysis is provided at GitHub (https://github.com/manuhuth/Data-Leakage-From-Covariances.git).

## Data availability

The CNSIM1 data set which we used in the manuscript is available within the DSData R package (https://github.com/datashield/DSData/tree/main/data).

## Acknowledgments

We thank the Interdisciplinary Research Unit Mathematics and Life Sciences at the University of Bonn, Nina Schmid, and Marc Vaisband for comments and discussions.

## Funding

This study was funded by the German Research Foundation (Deutsche Forschungsgemeinschaft, DFG) under Germany’s Excellence Strategy (EXC 2047 - 390873048 & EXC 2151 - 390685813), the German Ministry for Education and Research (Deutches Bundesminsterium für Bildung und Forschung, BMBF) under the CompLS program (grant agreement No 031L0293C), the University of Bonn (via the Schlegel Professorship of JH), the Helmholtz Association - Munich School for Data Science (MUDS), and the ORCHESTRA project. The ORCHESTRA project has received funding from the European Union’s Horizon 2020 research and innovation program under grant agreement No 101016167. The views expressed in this paper are the sole responsibility of the authors and the Commission is not responsible for any use that may be made of the information it contains. The funders had no role in the study design, data collection, data analyses, data interpretation, writing, or submission of this manuscript.

## Author contributions

M.H. developed the Covariance-Based Attack Algorithm. M.H., L.C. and J.H. proved the reconstruction accuracy. M.H. implemented the algorithm in DataSHIELD. R.G. implemented the algorithm in TensorFlow Federated. J.A. examined differential privacy and the tools Nvidia Flare, IBM federated learning, Intel’s OpenFL, and PySyft. J.H and E.T. conceptualized the study. M.H., J.A., and J.H. wrote the manuscript. All authors read and approved the final manuscript.

## Ethics declarations

The authors have no competing interests.

## Supplementary Figures

**Figure 6:**
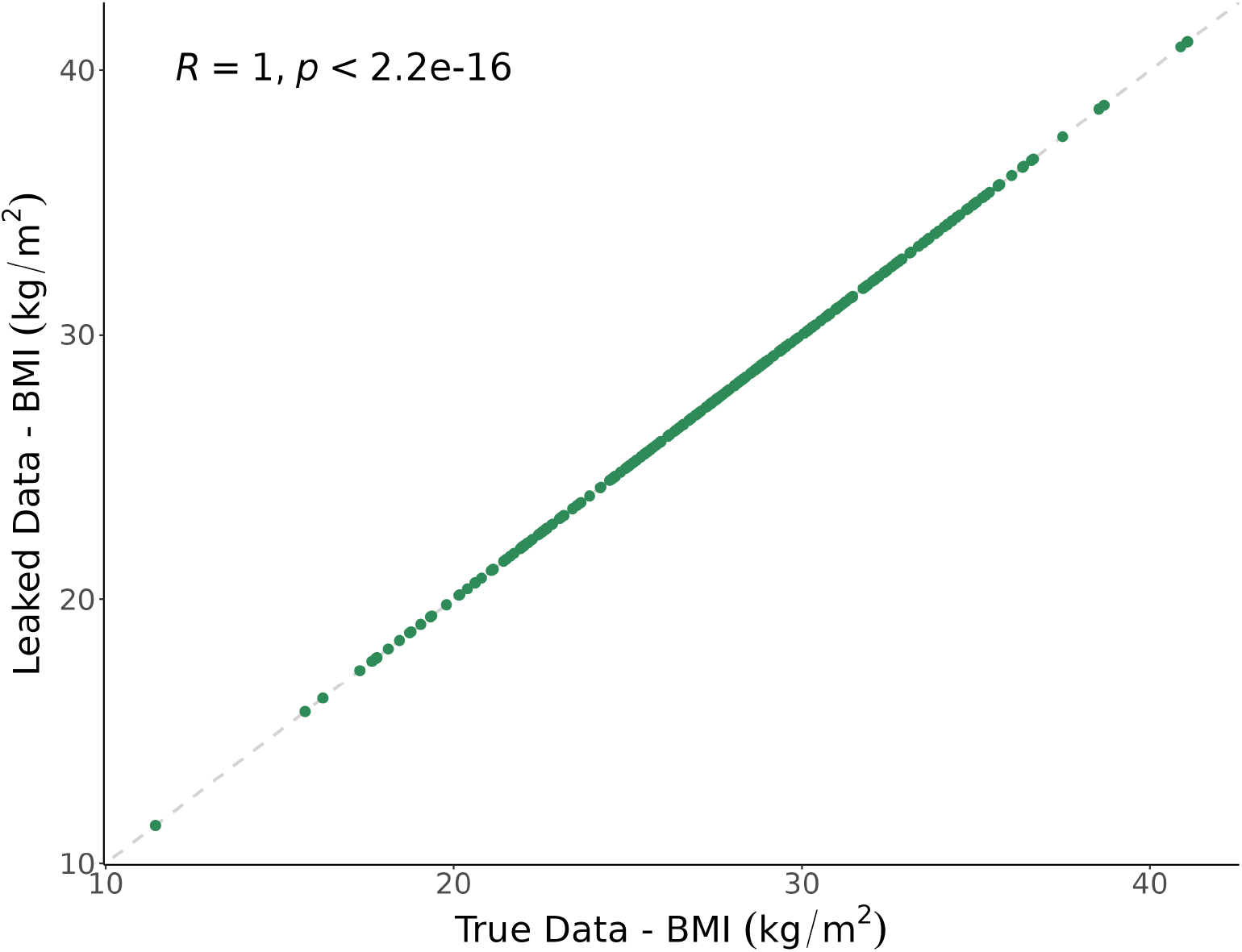
Leakage results for TensorFlow Federated are shown. The true data values from the first server of the CNSIM data set are plotted against the corresponding leaked data provided by the Covariance-Based Attack Algorithm.

## Footnotes

1 Note that the developers of TensorFlow Federated use the terms client and server in a way opposite to that of the DataSHIELD community. To avoid confusing the reader, we stick to the convention of DataSHIELD, with the client being the central hub and the servers being the data owners.

2 https://developer.nvidia.com/flare

## Notes

### Competing Interest Statement

The authors have declared no competing interest.

### Summary of Updates

We included an additional section analyzing differential privacy and further federated learning platforms.

